# Choline kinase A is required for alkylating agent resistance via lipid droplet formation

**DOI:** 10.64898/2026.11.05.764631

**Authors:** Abigail Fish, Kimberly Bodaar, Anais Barthe, Haven Griffin, Hasan Al Reza, Wojciech Rosikiewicz, Eric Aboagye, Julie R. Reisz, Angelo D’Alessandro, Chi-Lun Chang, Alejandro Gutierrez

**Author notes:** Equal contribution (Abigail Fish, Kimberly Bodaar). Corresponding author: Alejandro Gutierrez Department of Oncology St. Jude Children’s Research Hospital 262 Danny Thomas Place Memphis, TN 38105, USA.

## Abstract

Tolerance of DNA damage promotes cellular fitness, particularly in cancer cells under chemotherapeutic pressure. However, the biology and regulation of DNA damage responses remain incompletely understood. Using an unbiased genetic screen, we found that choline kinase A (CHKA) is required for alkylating resistance in T-cell acute lymphoblastic leukemia (T-ALL). CHKA is best-known as a cytoplasmic enzyme that catalyzes the first step in phosphatidylcholine biosynthesis, but alkylating agent resistance was mediated by a nuclear function of CHKA. We found that CHKA positively regulates transcription of the DNA repair factor RAD51 via formation of lipid droplets. Restoring lipid droplet formation or RAD51 expression was sufficient to confer alkylating agent resistance despite CHKA inactivation. Our findings link lipid droplet formation to transcriptional control of the DNA damage response, providing a mechanism through which cellular lipid metabolism can control the maintenance of genome integrity.

## INTRODUCTION

Alkylating agents are a class of reactive molecules ubiquitously encountered by living organisms, as natural products or pollutants in the environment and as products of endogenous cellular metabolism.^1^ These compounds are strong electrophiles that form covalent bonds with nucleophiles within biomolecules, with the ring nitrogen and extracyclic oxygen atoms of nucleotides particularly susceptible to alkylation.^2^ Thus, alkylating agent-induced DNA damage is a ubiquitous threat to genome integrity, and multiple molecular pathways have evolved that detect and repair the covalent DNA adducts that result.

The potential of pharmacologic alkylating agents as cancer therapeutics was initially suspected from the observation that the chemical warfare agent sulfur mustard (mustard gas) is profoundly toxic to hematopoietic cells.^3^ The subsequent testing of sulfur mustard analogs with improved pharmacologic properties, such as mechlorethamine (nitrogen mustard) and its many derivatives, revealed potent clinical activity in patients with hematopoietic and other tumor types.^4,5^ Several chemically distinct classes of alkylating agents have been developed as drugs, with nitrogen mustards such as mechlorethamine and cyclophosphamide comprising the largest class.^6^ Nitrogen mustards remain important components of contemporary treatment regimens for many cancers, including in front-line treatment regimens for acute lymphoblastic leukemia (ALL), and as intensive conditioning regimens for hematopoietic stem cell transplantation (HSCT) for relapsed/refractory leukemias.^7–10^ Prognosis for patients whose leukemias relapse after such HSCT regimens is extremely poor.^11–13^

Nitrogen mustards are bifunctional alkylating agents that can covalently bind to two distinct nucleophiles and often result in the formation of DNA interstrand cross-links. Multiple pathways have been linked to repair of DNA damage induced by these agents, including base excision repair, mismatch repair, homologous recombination, Fanconi Anemia and translesion synthesis pathways.^1^ While the cytotoxicity of alkylating agents is widely considered a direct consequence of DNA damage, metabolic state is a critical determinant of therapeutic response to these drugs.^14,15^ The molecular mechanisms through which cellular metabolism regulates the DNA damage response remain incompletely defined.

In this study, we used an unbiased genetic screen to find that choline kinase A (CHKA) is required for resistance to multiple alkylating agents, but not to other cytotoxics. CHKA is best-known as a cytoplasmic enzyme that catalyzes production of phosphocholine, the head group for the membrane phospholipid phosphatidylcholine. However, alkylating agent resistance was mediated by a nuclear function of CHKA. We found that CHKA is required for expression of the homologous recombination factor RAD51 via formation of lipid droplets. Indeed, restoring lipid droplet formation or RAD51 expression was sufficient to restore alkylating agent resistance following CHKA inactivation. Our data support the model that lipid droplets bind and sequester the transcriptional repressor MLX, and loss of lipid droplets leads to the MLX-dependent downregulation of RAD51. Our findings provide a molecular mechanism linking lipid metabolism to transcriptional control of the DNA damage response.

## RESULTS

### Choline Kinase A is required for alkylating agent resistance in T-cell acute lymphoblastic leukemia

To define molecular factors required for resistance to alkylating agents, we performed genome-wide CRISPR/Cas9 loss-of-function screens in the T-ALL cell lines Jurkat and PF382. Cas9-expressing T-ALL cells were transduced with the Brunello genome-wide CRISPR knockout library,^16^ treated with either vehicle or mechlorethamine, and guide RNA representation was assessed (Figure 1A). MAGeCK analysis was used to nominate genes required for alkylating agent resistance, on the basis of consistent depletion of guide RNAs targeting these genes upon mechlorethamine treatment in both cell lines. This analysis revealed genes that function in several pathways with well-established links to alkylating agent response, including the Fanconi pathway/interstrand cross-link repair, homologous recombination, nucleotide excision repair and glutathione biosynthesis (Figure 1B; Table S1).^1,17^ Among the top “hits” from this screen was choline kinase A (*CHKA*) (Figure 1B), which catalyzes the first step of the so-called Kennedy pathway for phosphatidylcholine biosynthesis.^18^ We prioritized CHKA for additional investigation because it is not an established component of the DNA damage response.

**Figure 1.**
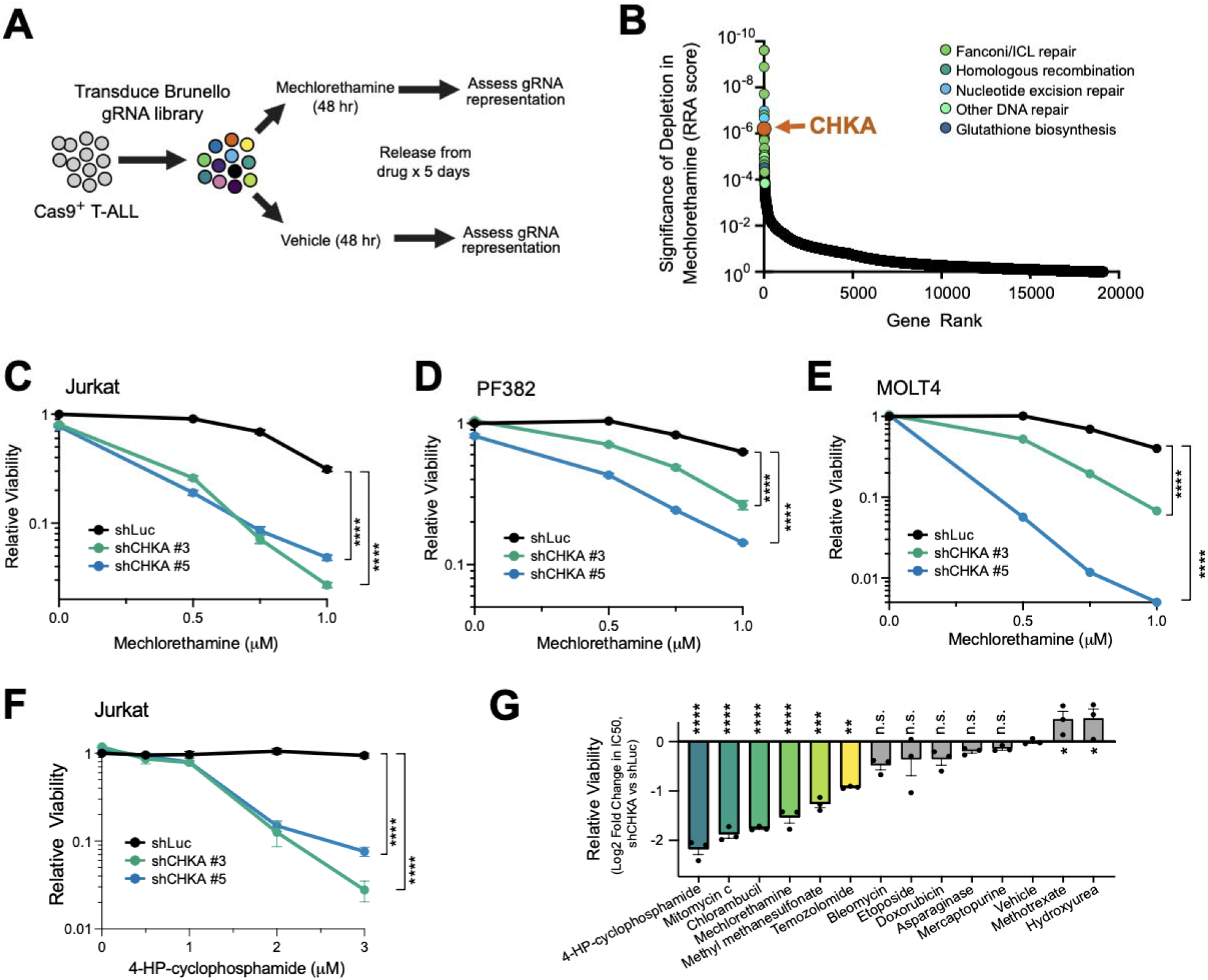
CHKA is required for alkylating agent resistance in T-ALL. (A) Experimental schema. (B) Results of the CRISPR/Cas9 screen, with genes ranked based on significance of depletion (RRA score) in mechlorethamine. Most of the top “hits” (n = 29 of 50) could be annotated into one of the functional categories shown. (C-F) The indicated T-ALL cells were transduced with shRNAs targeting CHKA or control (shLuciferase), treated with alkylating agents as indicated, and relative cell viability was assessed after 5 days of treatment. All viability results are normalized to those in shLuciferase, vehicle-treated controls. Significance was assessed at the highest alkylating agent dose shown using 2-way ANOVA with Dunnett adjustment for multiple comparisons. (G) Jurkat cells were transduced with shLuc or shCHKA #3, treated with vehicle or the indicated drugs at a series of doses that spanned the IC50 in control cells, and viability was assessed. Plot shows the ratio of IC50 values in CHKA-knockdown (shCHKA) versus control (shLuc) cells. Significance of differences versus vehicle-treated controls was assessed by one-way ANOVA with Dunnett adjustment for multiple comparisons. Results shown in (C-G) represent the mean ± SEM of at least n = 3 biological replicates. *, p < 0.05; **, p < 0.01; ***, p < 0.001; ****, p < 0.001; n.s., p ≥ 0.05.

To validate our screen while avoiding any potential interaction with CRISPR/Cas9-induced DNA damage, we turned to shRNA knockdown. We transduced 3 distinct T-ALL cell lines with shRNAs targeting *CHKA* or shLuciferase control, confirmed knockdown efficiency by Q-RT-PCR analysis, and found that CHKA knockdown induced sensitivity to the alkylating agents mechlorethamine and 4-hydroperoxycyclophosphamide (4-HP-cyclophosphamide; an active derivative of cyclophosphamide) (Figures 1C-F and S1A). Of note, mechlorethamine and 4-HP-cyclophophamide had indistinguishable effects on CHKA-dependent phenotypes assessed in our study, so these alkylating agents were used interchangeably throughout.

To distinguish whether CHKA is a specific regulator of alkylating agent response or a more general drug resistance factor, we transduced Jurkat cells with control or CHKA-targeting shRNAs, treated with a panel of cancer therapeutics, and assessed the degree to which CHKA knockdown altered drug sensitivity. This revealed that CHKA knockdown most strongly sensitized these cells to alkylating agents, including bifunctional (4-HP-cyclophosphamide, mitomycin C, chlorambucil, mechlorethamine) and monofunctional (methyl methanesulfonate, temozolomide) alkylators (Figure 1G). CHKA knockdown had a weak effect below the threshold of statistical significance on response to bleomycin, etoposide and doxorubicin, which induce DNA damage via free radical production (bleomycin, doxorubicin) and topoisomerase II inhibition (doxorubicin, etoposide).^2^ By contrast, CHKA knockdown did not induce sensitivity to drugs that disrupt homeostasis of nucleotides (mercaptopurine, methotrexate, hydroxyurea), amino acids (asparaginase), or microtubule polymerization (vincristine) (Figure 1G). To test whether CHKA function is differentially required for alkylating agent response in malignant versus normal hematopoietic cells, we then transduced CD34+ hematopoietic stem and progenitor cells (HSPCs) collected from healthy human donors with these same shRNAs and treated cells with mechlorethamine. We found that *CHKA* knockdown had no significant effect on the ability of normal HSPCs to tolerate mechlorethamine (Figures S1B-C).

### The kinase activity of CHKA is required for alkylating agent resistance

To test whether the enzymatic activity of CHKA is required for drug resistance, we tested whether the toxicity of CHKA knockdown could be reversed by restoring expression of wild-type CHKA, or of a catalytically defective D306A point mutant.^19,20^ Jurkat cells were first transduced with cDNA expression constructs encoding these CHKA alleles with wobble-base substitutions to escape shRNA targeting (versus EGFP control), and then with shRNAs targeting endogenous CHKA or shLuciferase control. We confirmed that transduction of these shRNAs and cDNAs had the expected effects on CHKA protein expression by Western blot analysis (Figure S2). Cells were then treated with mechlorethamine, and viability was assessed. We found that the toxicity of CHKA depletion was reversed by expression of wild-type CHKA, but not the catalytically dead CHKA allele (Figure 2A). We also treated cells with the enzymatic CHKA inhibitors CK2 (also known as ICL-CCIC-0019) or CK14,^21,22^ and found that these phenocopied alkylating agent sensitivity in Jurkat cells and in patient-derived models of T-ALL (Figures 2B-C). Thus, alkylating agent resistance requires the kinase activity of CHKA.

**Figure 2.**
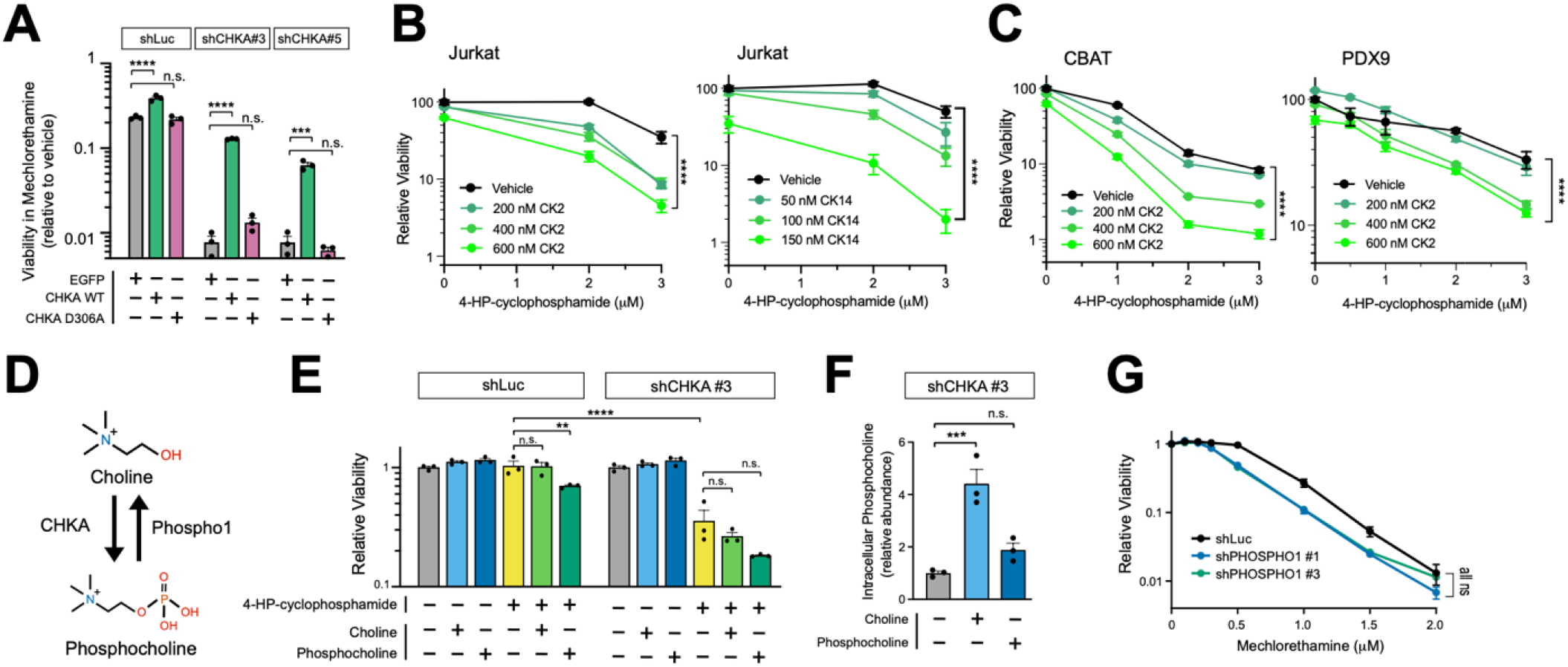
The enzymatic activity of CHKA is required for alkylating agent resistance. (A) Jurkat T-ALL cells were transduced with the indicated shRNAs together with cDNA rescue constructs encoding EGFP (negative control) or the indicated CHKA alleles. Cell viability was assessed after 5 days of treatment with 1 µM mechlorethamine using CellTiter-Glo analysis. Cell viability in all conditions was normalized to that in vehicle-treated, shLuciferase and EGFP-transduced controls. Significance was assessed by 2-way ANOVA with Dunnett adjustment for multiple comparisons. (B) and (C) Jurkat cells (B) or the indicated patient-derived xenograft models (C) were pre-treated overnight with the substrate-competitive CHKA inhibitors CK2 or CK14, then with 4-HP-cyclophosphamide at the indicated doses for 4 days, and cell viability was assessed as in (A). Significance was assessed by 2-way ANOVA using the highest dose of 4-HP-cyclophosphamide. (D) Schematic of canonical CHKA and PHOSPHO1 functions. (E) Jurkat T-ALL cells were transduced with the indicated shRNAs and supplemented with the indicated metabolites (1 mM added daily) beginning 24 hours before starting treatment with 4-HP-cyclophosphamide. Relative cell viability was assessed after 5 days of treatment as in (A). A two-way ANOVA with a Sidak correction was used for statistical analysis. (F) Jurkat cells transduced with shCHKA #3 were supplemented with the indicated metabolite for 24 hours, and intracellular phosphocholine abundance was assessed by metabolomics analysis. Significance was assessed by one-way ANOVA with Dunnett adjustment for multiple comparisons. (G) Jurkat cells transduced with the indicated shRNAs were treated with mechlorethamine at the indicated doses, and viability was assessed after 5 days of treatment. Significance was assessed by Welch’s t test. Results shown in (A-G) represent the mean ± SEM of at least n = 3 biological replicates. *, p < 0.05; **, p < 0.01; ***, p < 0.001; ****, p < 0.001; n.s., p ≥ 0.05.

The best-known function of CHKA is production of phosphocholine (Figure 2D).^23,24^ If alkylating agent resistance is dependent on this function of CHKA, we reasoned that supplementing CHKA knockdown cells with phosphocholine might rescue this phenotype. However, we found that adding phosphocholine to the media did not protect CHKA-knockdown cells from the toxicity of 4-HP-cyclophosphamide (Figure 2E), although experimental interpretation was complicated by the very modest increase in intracellular phosphocholine levels achieved (Figure 2F), reflecting the poor membrane permeability of phosphocholine. We also tested choline supplementation, reasoning that i) this molecule ought to have more favorable bioavailability due to established cell membrane transporters,^23^ and ii) because shRNA knockdown reduced but did not eliminate expression of the CHKA enzyme (Figure S2), increasing substrate concentration might effectively increase phosphocholine production in these cells. While choline supplementation did significantly increase intracellular phosphocholine concentrations in CHKA-knockdown cells, this did not protect these cells from alkylating agent toxicity (Figures 2E-F). If CHKA knockdown induces alkylating agent sensitivity by impairing phosphocholine production, we also reasoned that knockdown of the small molecule phosphatase PHOSPHO1, which antagonizes this canonical function of CHKA (Figure 2D),^25^ might be expected to increase alkylating agent resistance, but this was not the case (Figures 2G and S3A). CHKA also functions as a lower-affinity kinase for ethanolamine,^24^ but phosphoethanolamine supplementation did not protect CHKA-knockdown cells from alkylating agent toxicity (Figure S3B), and knockdown of *CHKB*, the higher-affinity catalyst of phosphoethanolamine production,^24^ did not phenocopy alkylating agent sensitivity (Figures S3C-D). To further probe whether other choline/phosphocholine related metabolites influenced alkylating agent sensitivity, we also tested supplementation with betaine, folate, S-adenosylmethionine or phosphatidylcholine. However, none of these metabolites were able to protect CHKA-knockdown cells from the toxicity of alkylating agents (Figure S4).

### Alkylating agent resistance is mediated by a nuclear function of CHKA

The canonical function of CHKA is cytoplasmic,^23^ but the above data suggested that alkylating agent resistance might reflect an alternative function of CHKA. To investigate this possibility, we began by assessing subcellular localization of the CHKA protein and found that it localized to the nucleus as well as the cytoplasm (Figure 3A). To distinguish whether alkylating agent resistance was mediated by a nuclear or a cytoplasmic function of CHKA, we generated lentiviral expression constructs encoding wild-type CHKA fused to an N-terminal HA-tag and nuclear localization signal (HA-NLS) or an HA-tag nuclear export signal (HA-NES). These were transduced into Jurkat cells, and we confirmed appropriate localization of each fusion protein (Figure 3B). Cells transduced with these constructs (or CHKA wild-type, the catalytically impaired CHKA-D306A, or EGFP as additional controls) were then transduced with shRNAs targeting endogenous CHKA or shLuciferase control. Note that all CHKA cDNAs utilized contained wobble-base substitutions to escape shRNA targeting, and we validated that these perturbations altered CHKA protein levels as expected (Figure S5). These cells were then treated with 4-HP-cyclophosphamide, and viability was assessed. We found that the nuclear-localized CHKA was able to restore alkylating agent resistance at least as well as wild-type CHKA, whereas cytoplasmically localized CHKA was as functionally impaired as the catalytically dead D306A mutant (Figure 3C). These findings indicate that alkylating agent resistance is mediated by a nuclear function of CHKA.

**Figure 3.**
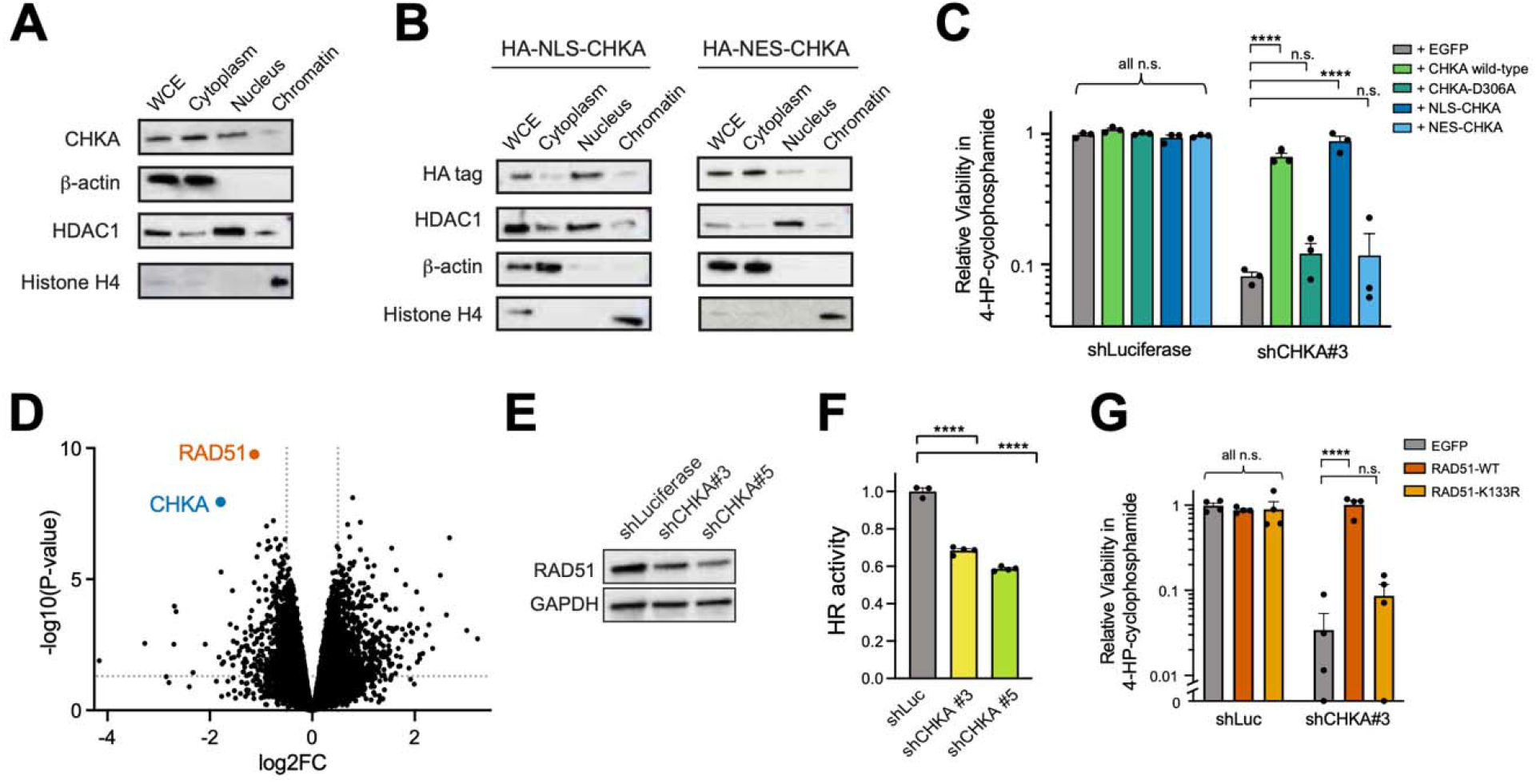
Nuclear CHKA regulates alkylating agent resistance via transcriptional regulation of *RAD51.* (A) Jurkat cell lysates were subjected to subcellular fractionation by centrifugation, and fractions were analyzed by Western blot for the indicated proteins. (B) Jurkat cells transduced with the indicated expression constructs were subjected to subcellular fractionation and analyzed by Western blot for the indicated proteins. (C) Jurkat T-ALL cells were transduced with the indicated shRNAs together with cDNA rescue constructs encoding EGFP (negative control) or the indicated CHKA alleles, treated for 5 days with 2 µM 4-HP Cyclophosphamide, and viability was assessed by CellTiter-Glo analysis. Viability of all cells is normalized to that in shLuciferase controls. Significance was assessed using 2-way ANOVA with Dunnett adjustment for multiple comparisons. (D) Jurkat were transduced with CHKA or Luciferase-targeting shRNAs, and RNA-sequencing differential expression analysis was performed. (E) Jurkat cells were transduced with the indicated shRNAs, and expression of the indicated proteins was assessed by Western blot analysis. (F) Jurkat cells transduced with the indicated shRNAs were analyzed for homologous recombination activity based on repair of co-transduced plasmids. Results are normalized to those in shLuciferase-transduced controls. An Ordinary One-Way ANOVA test was used to verify statistical significance. (G) Jurkat T-ALL cells were transduced with the indicated shRNAs together with cDNA rescue constructs encoding EGFP (negative control) or the indicated RAD51 alleles. Cells were treated with 2 µM 4-HP-cyclophosphamide for 5 days, and relative cell viability was assessed using CellTiter-Glo analysis. Significance was assessed using 2-way ANOVA with Dunnett adjustment for multiple comparisons. Results shown in (C, F-G) represent the mean ± SEM of at least n = 3 biological replicates. *, p < 0.05; **, p < 0.01; ***, p < 0.001; ****, p < 0.001; n.s., p ≥ 0.05.

We then asked how nuclear CHKA regulates alkylating agent response. Using RNA-seq and ATAC-seq analysis, we found that CHKA knockdown altered mRNA expression and chromatin accessibility at a number of loci (Figures 3D and S6A; Tables S2 and S3). Strikingly, the single most significantly downregulated transcript following CHKA knockdown was *RAD51* (Figure 3D), which was also downregulated at the protein level (Figure 3E). *RAD51* encodes a recombinase that catalyzes strand exchange during homologous recombination (HR),^26,27^ whose inhibition increases alkylating agent sensitivity in multiple contexts.^28–30^ Using a two-plasmid reporter system designed to measure HR efficiency, we found that CHKA knockdown impaired homologous recombination (Figure 3F). We then asked whether transcriptional downregulation of RAD51 causes alkylating agent sensitivity following CHKA knockdown. Jurkat cells were first transduced with cDNA expression constructs encoding wild-type RAD51, a RAD51-K133R ATPase-defective mutant^31^ or EGFP control, and subsequently with shRNAs targeting CHKA or shLuciferase control. Cells were then treated with 4-HP-cyclophosphamide, and viability was assessed. We found that the toxicity of CHKA knockdown was completely rescued by restoring expression of wild-type RAD51, but not the K133R ATPase-defective mutant. From these data, we conclude that CHKA loss increases the toxicity of alkylating agents via transcriptional downregulation of the homologous recombination factor RAD51.

### CHKA controls RAD51 expression and alkylating agent response via formation of lipid droplets

The above data supported the model that a nuclear enzymatic function of CHKA controls RAD51 transcription and alkylating agent response. However, CHKA is not an established transcriptional regulator, and we found no evidence of homology between defined or unannotated regions of the CHKA protein and any established DNA-binding, transcriptional or chromatin-regulatory domains using either sequence-based (HMMER,^32^ BLAST^33^) or structure-based (Foldseek^34^) approaches. In considering possible mechanisms through which CHKA might regulate gene expression indirectly, our attention was drawn to lipid droplets because loss of multiple phosphatidylcholine biosynthetic factors, including the CHKA ortholog in Drosophila, disrupts lipid droplet homeostasis in flies and mammals.^35,36^ Lipid droplets are cellular organelles comprised of a core of neutral lipids (triglycerides, sterol esters) enclosed by a phosphatidylcholine-rich phospholipid monolayer, with several associated proteins.^37–39^ While best-known for their roles in energy storage and cellular metabolism, lipid droplets can also regulate gene expression by binding and sequestering some transcription factors,^40,41^ providing a potential link between CHKA function and transcriptional regulation of RAD51.

To test whether CHKA regulates lipid droplet formation, we transduced Jurkat cells with control or CHKA-targeting shRNAs and used fluorescence microscopy analysis after staining with the lipid marker BODIPY 493/503 to quantify lipid droplet formation. This revealed that CHKA knockdown impaired the formation of lipid droplets (Figures 4A-B). We then asked whether stimulating lipid droplet formation might protect CHKA-deficient cells from the toxicity of alkylating agent. Cells were transduced with control or CHKA-targeting shRNAs, and supplemented with vehicle or oleic acid, which stimulates lipid droplet formation.^35^ We confirmed that oleic acid treatment effectively stimulates lipid droplet formation in CHKA-depleted cells (Figure 4C), and this was sufficient to restore RAD51 expression and alkylating agent resistance in CHKA-deficient cells (Figures 4D-E). Moreover, treatment with a small molecule inhibitor of phospholipase A2α that impairs lipid droplet formation^42^ induced alkylating agent sensitivity (Figure 4F). These findings support the conclusion that lipid droplet formation is necessary and sufficient for RAD51 expression and alkylating agent resistance downstream of CHKA.

**Figure 4.**
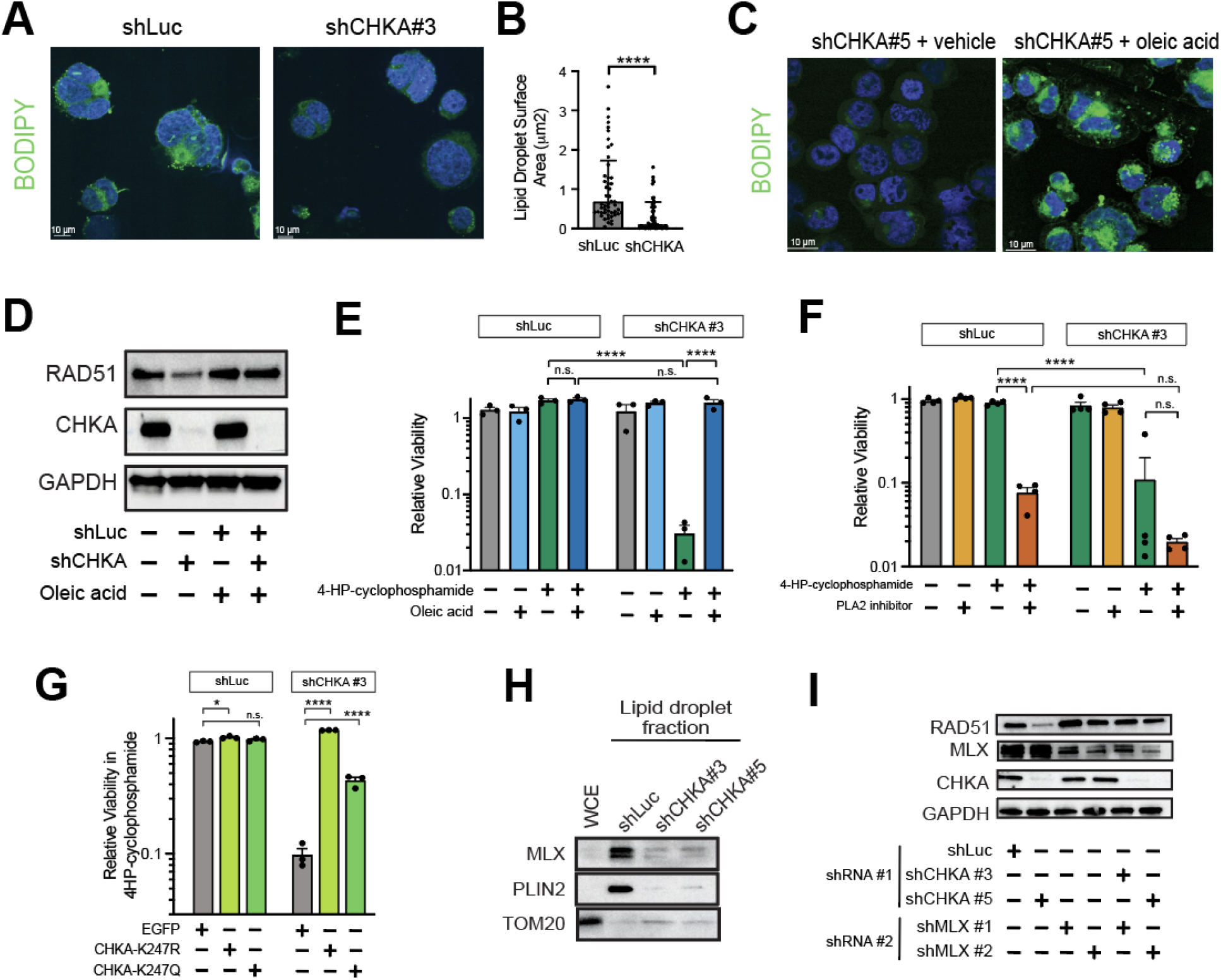
CHKA controls RAD51 expression and alkylating agent response via lipid droplets. (A) Jurkat cells transduced with the indicated shRNAs were imaged via fluorescence using BODIPY 488/503 (green) and DAPI (blue). Image brightness/contrast was adjusted uniformly across both panels to increase visibility. (B) Quantification of lipid droplet surface area per cell section in cells from (A). Fifty cells were quantified per condition. (C) Jurkat cells were treated with 1 mM oleic acid or vehicle and imaged via fluorescence using BODIPY 488/503 (green) and DAPI (blue) staining. Image brightness/contrast was adjusted uniformly across both panels to increase visibility. A Welch’s t-test was used for statistical analysis. (D) Jurkat T-ALL cells were transduced with the indicated shRNAs, and Western blot analysis was performed for the indicated proteins. (E) Jurkat T-ALL were transduced with the indicated shRNAs and, where indicated, supplemented with 1mM oleic acid 24 hours before starting treatment with 4-HP-cyclophosphamide and every 24 hours thereafter for five days. Relative cell viability was assessed after 5 days of treatment with 2 µM 4-HP-cyclophosphamide using CellTiter-Glo analysis. Significance was assessed by 2-way ANOVA with Tukey’s adjustment for multiple comparisons. (F) Jurkat T-ALL cells were transduced with the indicated shRNAs and supplemented with 15 nM of cytosolic phospholipase A2α inhibitor CAY10650 24 hours before starting treatment with 4-HP-cyclophosphamide and every 24 hours thereafter for five days. Relative cell viability was assessed after 5 days of treatment with 2 µM 4-HP-cyclophosphamide using CellTiter-Glo analysis. Significance was assessed by 2-way ANOVA with Tukey’s adjustment for multiple comparisons. (G) Jurkat T-ALL cells were transduced with the indicated shRNAs together with cDNA rescue constructs encoding EGFP (negative control) or the indicated CHKA K247 alleles. Cells were treated with 2 µM 4-HP-cyclophosphamide for 5 days, and relative cell viability was assessed using CellTiter-Glo analysis. Significance was assessed using 2-way ANOVA with Dunnett adjustment for multiple comparisons. (H) Lipid droplets were extracted from 15 million cells per condition, using Jurkat cells transduced with the indicated shRNAs. The whole cell extract (WCE) is from shLuciferase controls. 5.5% of the lipid droplet fraction of each sample was loaded on the gel and analyzed by Western blot analysis for the indicated proteins. (I) Jurkat cells were transduced with the indicated shRNAs, and expression of the indicated proteins was assessed by Western blot analysis. Results shown in (E-G) represent the mean ± SEM of at least n = 3 biological replicates. *, p < 0.05; **, p < 0.01; ***, p < 0.001; ****, p < 0.001; n.s., p ≥ 0.05.

**Figure 5.**
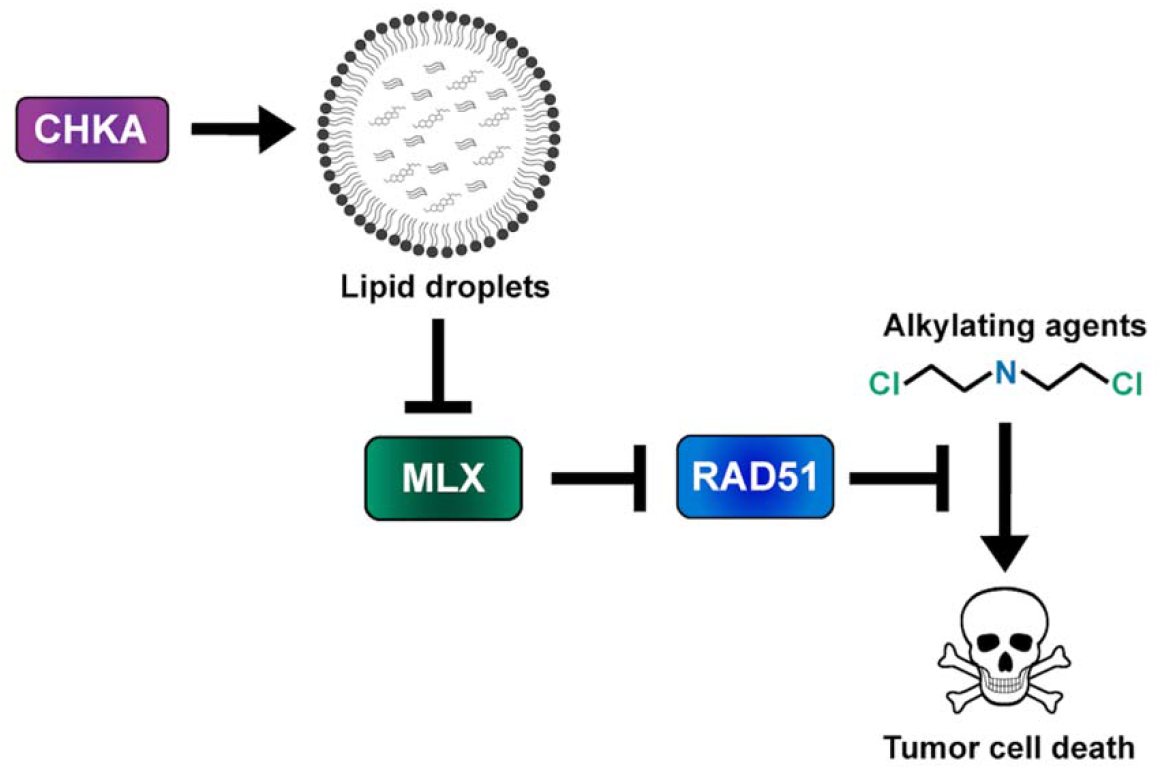
Proposed model.

In addition to the requirement for CHKA and other phosphatidylcholine biosynthetic factors for lipid droplet formation,^35,36^ CHKA has also been shown to bind to lipid droplets, phosphorylate PLIN2/3 and promote lipolysis in a manner dependent on acetylation of CHKA lysine 247.^43^ To test whether the role of CHKA in alkylating agent response might reflect this alternative function of CHKA, we leverage an acetylation-dead CHKA-K247R mutant whose ability to bind and phosphorylate PLIN2/3 and initiate lipolysis is impaired. Transduction of CHKA-depleted cells with a CHKA-K247R expression construct that escapes shRNA targeting fully restored alkylating agent resistance (Figure 4G), whereas the acetylation-mimetic CHKA-K247Q had a weaker effect (Figure 4G). These findings suggest that the role of CHKA in alkylating agent resistance is independent of its ability to bind lipid droplets and stimulate lipolysis.

We then asked how lipid droplet formation regulates RAD51 expression. Lipid droplets have been shown to bind and sequester the transcription factor MLX.^40^ This finding caught our attention because our ATAC-seq analysis revealed that many of the genomic loci whose chromatin accessibility decreased following CHKA knockdown are MYC target genes (Figure S6B; Table S4), and MLX can repress transcription at canonical MYC E-box motifs.^44–46^ We isolated lipid droplets by differential centrifugation and confirmed that MLX is bound to lipid droplets in control (shLuciferase-transduced) Jurkat cells (Figure 4H). CHKA knockdown markedly reduced levels of both MLX and of the lipid droplet protein PLIN2 from the lipid droplet fraction of these cells (Figure 4H), without changing total MLX protein levels in these cells (Figure 4I, left two lanes). This suggested reduced sequestration of MLX by lipid droplets as a consequence of reduced lipid droplet content in CHKA-depleted cells. We then asked whether MLX is required for repression of RAD51 in CHKA-knockdown cells. Indeed, we found that MLX knockdown prevented the downregulation of RAD51 by CHKA knockdown (Figure 4I).

## DISCUSSION

Our work has revealed that homologous recombination and alkylating agent resistance are unexpectedly regulated by lipid droplets, a cellular organelle best-known for its roles in energy storage and cellular metabolism. Using a genome-wide CRISPR/Cas9 screen, we found that CHKA is required for resistance to multiple alkylating agents, in a manner dependent on its kinase activity and nuclear localization. CHKA depletion reduced expression of RAD51, impaired homologous recombination, and increased sensitivity to alkylating agents. Notably, despite the broad transcriptional consequences of CHKA loss, restoring expression of RAD51 alone was sufficient to restore alkylating agent resistance. These findings establish RAD51 downregulation as a key functional consequence of CHKA loss that leads to alkylating agent sensitivity. The finding that CHKA depletion selectively increased sensitivity to alkylating agents, with substantially weaker effects on other classes of cytotoxics, indicates a specific dependence on this pathway for the tolerance of alkylating agent-induced DNA damage, rather than a more general requirement for CHKA in the tolerance of cellular stress.

Our findings connect this DNA repair phenotype to lipid droplet biology. Previous work has shown that lipid droplets regulate the expression of metabolic gene expression programs by binding and sequestering the transcription factor MLX.^40^ Our findings extend this function of lipid droplets to the control of a core DNA repair factor. Indeed, CHKA depletion impaired lipid droplet formation, reduced association of MLX with the lipid droplet fraction, and resulted in the MLX-dependent repression of RAD51. Moreover, stimulating lipid droplet formation with oleic acid was sufficient to restore RAD51 expression and alkylating agent resistance in CHKA-deficient cells. These findings place lipid droplets upstream of RAD51, and suggest that the physiologic state of this organelle, best-known for its role in energy storage and cellular metabolism, controls the cellular response to DNA damage.

More broadly, our findings suggest that lipid droplets provide a mechanism for coordinating genome maintenance with cellular metabolic state. Homologous recombination functions principally during the S and G2 phases of the cell cycle, when DNA lesions can be repaired using an intact sister chromatid. Regulation of RAD51 expression by lipid droplet homeostasis could therefore provide a mechanism to coordinate DNA repair capacity with the metabolic programs that support cell growth and proliferation. However, lipid droplets are dynamic organelles whose composition and function change in response to diverse metabolic and cellular cues, and their relationship to nutrient availability is complex. Thus, it remains unclear which physiological perturbations of lipid droplet homeostasis might engage this pathway. Understanding these relationships is expected to reveal additional mechanisms through which cellular metabolism regulates the maintenance of genome integrity.

## MATERIALS AND METHODS

### Cell Lines and Cell Culture

The 293T cell line was obtained from ATCC (Manassas, VA, USA) and cultured in DMEM (Thermo Fisher Scientific) with 10% fetal bovine serum (Life Technologies Corporation, Carlsbad, CA) and 1% penicillin/streptomycin (Thermo Fisher Scientific) at 37°C, 5% CO2. T-ALL and AML cell lines were obtained from ATCC (Manassas, VA, USA) or DSMZ (Braunschweig, Germany) and cultured in RPMI-1640 (Thermo Fisher Scientific) with 10% FBS and 1% penicillin/streptomycin. Human CD34+ hematopoietic stem and progenitor cells (HSPCs) from mobilized peripheral blood of healthy donors were obtained from Fred Hutchinson Cancer Research Center (Seattle, WA, USA). HSPCs were cultured in IMDM (Thermo Fisher Scientific) supplemented with 20% FBS and recombinant human interleukin-3 (R&D systems, Minneapolis, MN), recombinant human interleukin-6 (R&D systems) and recombinant human stem cell factor (R&D systems) to a final concentration of 50 ng/ml each.

Cell line identities were validated using short tandem repeat profiling at Dana-Farber Cancer Institute or St. Jude Children’s Research Hospital core facilities, at which point multiple vials were frozen and freshly verified stocks were used for all experiments. Mycoplasma contamination was excluded routinely throughout the course of these experiments using the MycoAlert Mycoplasma Detection Kit (Lonza, Basel, Switzerland).

Patient-Derived Xenograft (PDX) samples were obtained from the Dana-Farber Cancer Institute Public Repository of Xenografts,^47^ and were cultured in alpha-minimum essential media supplemented with 10% fetal bovine serum (Life Technologies Corporation), 10% human AB serum (Invitrogen, Carlsbad, CA), 1% L-glutamine (Thermo Fisher Scientific), 1% penicillin/streptomycin (Thermo Fisher Scientific) in the presence of recombinant human cytokines stem cell factor (50 ng/mL), Flt3-L (20 ng/mL) and IL-7 (10 ng/mL)(R&D Systems) at 37°C under 5% CO2, as previously described.^48^ PDX samples were thawed and incubated overnight before plating for experiments to select out death cells from freeze-thawing.

### Genome-Wide Loss of Function Screen

The genome-wide loss-of function screen was performed using the human Brunello CRISPR knockout pooled library (a gift from David Root and John Doench; Addgene # 73178), as described.^16^ Jurkat and PF382 cells were first transduced with lentiCas9-blast (a gift from Feng Zhang; Addgene plasmid # 52962), selected with blasticidin, and Cas9 activity was confirmed using a self-excising GFP construct, pXPR_011 (a gift from John Doench and David Root; Addgene plasmid # 59702). The Brunello library was transduced in biologic duplicates per cell line into Cas9-expressing cells at a goal multiplicity of infection (MOI) < 0.3 (MOI achieved: PF382, 0.26; Jurkat, 0.20). Cells were split every other day and selected with puromycin (1 μg/ml) beginning 24 hours post transduction, which was continued for 8 days. Cells were treated with mechlorethamine at doses targeting 75% fractional inhibition (PF382, 0.9 µM; Jurkat, 2.5 μM; fractional inhibition achieved: PF382, 83%; Jurkat, 70%) for 48 hours beginning at day 10 post-infection with the Brunello library, and then released from the drug for 5 days, whereafter cells were harvested. The coverage achieved per Brunello gRNA was estimated at 475x. Genomic DNA was extracted using the Blood & Cell Culture DNA Maxi Kit (Qiagen). Guide RNA sequences were PCR-amplified from genomic DNA and quantified by sequencing using an Illumina NextSeq 500. Significance of gene depletion was assessed, based on selective depletion of guide RNAs targeting each gene in mechlorethamine-treated cells, using MAGeCK analysis of both cell lines combined (https://sourceforge.net/projects/mageck/).^49^

### shRNA and cDNA Expression Plasmids

Gateway Recombination Cloning Technology (Invitrogen) was used according to the manufacturer’s instructions to transfer in vitro synthesized gene strands (Twist Bioscience, San Francisco, CA) flanked by AttB sites, into the pDONR221 vector (Invitrogen #12536017) and then into pLenti CMV Neo Dest (Addgene #17392). This approach was used to generate expression constructs encoding codon-optimized wild-type CHKA, CHKA D306A, CHKA K247R, or CHKA K247Q (sequence based on NCBI CCDS8178), with wobble-base substitutions also included in the shRNA target sites to avoid targeting by CHKA-targeting shRNAs. Correct generation of each plasmid was confirmed by Sanger sequencing (Genewiz). The EGFP control expression plasmid was generated in the same manner. HA-NES-CHKA and HA-NLS-CHKA were generated by introducing the following sequences between the initiator ATG and the remainder of the CHKA protein-coding sequence into the previously generated wild-type CHKA expression plasmid in pLenti CMV Neo Dest, with cloning performed by Custom DNA Constructs LLC: HA-NES, gcctacccctacgacgtgcccgactacgcctccctcggatctctgccgccgctggagcgcctgacgctgggatcc; HA-NLS, gcctacccctacgacgtgcccgactacgcctccctcggatctctgccgccgctggagcgcctgacgctgggatcc.

The wild-type RAD51 or RAD51 K133R cDNA expression constructs (Supplemental Table 5) were generated by ATUM.bio in the pD2108-CMV lentiviral expression construct. The following lentiviral shRNA vectors in pLKO.1 with puromycin or neomycin resistance were generated by the RNAi Consortium library of the Broad Institute and obtained from Sigma-Aldrich: shLuciferase (TRCN0000072243), shCHKA #3 (TRCN0000006048), shCHKA #5 (TRCN0000006050), shCHKB #1 TRCN0000052553, shCHKB #4 TRCN0000052556, shPHOSPHO1 #1 (TRCN0000011059), shPHOSPHO1 #3 (TRCN0000006969), shMLX #1 (TRCN0000015722), shMLX #2 (TRCN0000015720), and shGFP (TRCN0000072181).

### Lentiviral Production, Infection and Selection

Lentiviruses were generated by co-transfecting lentiviral plasmids of interest together with packaging vectors psPAX2 (a gift from Didier Trono; Addgene plasmid # 12260) and VSV.G (a gift from Tannishtha Reya, Addgene plasmid # 14888) using polyethyleneimine, M.W. 25,000, linear (VWR International, Radnor, PA) in 293T cells. Lentiviral infection was performed by the spinoculation of cell lines with virus-containing media (1,500 g x 90 minutes) in the presence of 8 μg/ml polybrene (Merck Millipore, Darmstadt, Germany). Selection with puromycin (1 µg/ml for a minimum of 48 hours; Thermo Fisher Scientific), neomycin (750 µg/ml for a minimum of 6 days; Thermo Fisher Scientific) or blasticidin (15 μg/ml for a minimum of 5 days; Invitrogen) was begun 24 hours after infection. After selection, infected cells were used for the experiments described below. For all cDNA rescue experiments, sequential transduction of lentivirus plasmids was performed beginning with the neomycin or blasticidin resistant cDNA expression plasmid and followed by the puromycin-resistant shRNA plasmid.

### Compounds

Choline, acetylcholine, phosphocholine, doxorubicin, etoposide, mitomycin C, bleomycin, oleic acid, phosphoethanolamine, and mechlorethamine-chloride (nitrogen mustard) were obtained from Sigma Aldrich (St. Louis, MO, USA). 6-Mercaptopurine was obtained from Abcam (Cambridge, United Kingdom). The PEGylated asparaginase used (pegaspargase) was from Servier (Suresnes, France). Vincristine and methotrexate were obtained from Selleckchem (Houston, TX, USA). CK2 (ICL-CCIC-0019) and CK14 were a kind gift from Prof. Aboagye (Imperial College London).^22^ 4-HP-cyclophosphamide, cytosolic phospholipase A_2_α (cPLA_2_α) inhibitor (CAY10650), temozolamide, chlorambucil, S-adenosylmethionine, betaine, folic acid, 5-MTHF, phosphatidylcholine, hydroxyurea, and bleomycin were obtained from Cayman Chemical (Ann Arbor, MI, USA). Methyl methanesulfonate was from Sigma-Aldrich (#129925).

### Metabolomics

Jurkat cells transduced with shRNAs targeting CHKA or shLuc negative control were plated 1 million/mL and treated with vehicle (DMSO) or 5 mM choline/phosphocholine (Sigma Aldrich) for 24 hours in triplicate. Cells were washed 3 times with PBS, all PBS removed, then cells frozen as pellets and shipped on dry ice to the University of Colorado. Frozen pellets were placed on dry ice and ice cold lysis buffer (5:3:2 methanol:acetonitrile:water, all Optima grade from Fisher) was added at a concentration of 2,000,000 cells/mL. Samples were vortexed for 30 minutes at 4°C and centrifuged for 10 minutes at 18,213 g at 4°C. Supernatant was transferred to autosampler vials and analyzed (10 µL injection) using a Thermo Vanquish UHPLC coupled to a Thermo Q Exactive mass spectrometer using a 5-minute C18 gradient in positive and negative ion modes (separate runs) as previously described in detail.^50^ Following data acquisition, .raw files were converted to .mzXML using RawConverter then metabolites assigned and peaks integrated using Maven (Princeton University) in conjunction with the KEGG database and an in-house standard library. Quality control was assessed using technical replicates run at beginning, end, and middle of each sequence as previously described.^51,52^

### RNA Extraction and Quantitative RT-PCR Analysis

Following transduction with lentivirally-delivered shRNAs, mRNA expression was assessed by quantitative reverse transcriptase polymerase chain reaction (Q-RT-PCR). Total RNA was extracted using the RNeasy Mini Kit (Qiagen) according to the manufacturer’s instructions. cDNA was synthesized using the SuperScript IV First-Strand Synthesis system (Invitrogen) according to the manufacturer’s instructions. Q-RT-PCR analysis was performed with the SYBR Green PCR Core Reagents kit (Applied Biosystems, Life Technologies, Grand Island, NY, USA) and a 7500 Real Time PCR System instrument (Applied Biosystems) according to the manufacturer’s instructions. All Q-RT-PCR reactions were performed in triplicate using primers listed below. Cycle threshold (CT) for each condition, in triplicate, was compared to β-actin control according to the 2ΔCT comparative method.

Primers used were as follows (all 5′–3′):

Human β-actin Fwd AGAAAATCTGGCACCACACC

Human β-actin Rev AGAGGCGTACAGGGATAGCA

Human CHKA Fwd GTTTGCCAGATATTTCTGCAG

Human CHKA Rev CCAAGGGCAGATTGTAACTG

Human Phospho1 Fwd GAAGGGAGATTCGGCAAAGA

Human Phospho1 Rev CCGAGGTGGGTTAACTGAATAG

### Cell Viability Assays

10,000 cells/well were plated in 96-well plates and treated for 5 days with the indicated doses of mechlorethamine or 4-HP-cyclophosphamide, an active metabolite of cyclophosphamide. For metabolite rescue experiments, metabolites were added to the media at the indicated doses 24 hours before treatment started. Cell viability was assessed using CellTiter-Glo analysis (Promega). Results are normalized to vehicle control. For the testing of a multiple conventional chemotherapeutics (Figure 1G), 10,000 Jurkat cells/well were plated and treated with drug concentrations across a serial 10-fold dilution range spanning the IC50 concentrations, each in triplicate. Cell viability was assessed using CellTiter-Glo (Promega) analysis after 5 days and the Log2 Fold Change in IC50 of shCHKA/shLuc was graphed.

### Western Blot

Cells were lysed in RIPA buffer (Merck Millipore) supplemented with cOmplete protease inhibitor (Roche, Basel, Switzerland) and PhosSTOP phosphatase inhibitor (Roche). Thirty μg of protein lysate was mixed with Laemmli sample buffer (Bio-Rad, Hercules, CA) and β-mercaptoethanol (Sigma-Aldrich) before being run on a 4–12% Novex bis-tris polyacrylamide gel (Thermo Fisher Scientific) or 4-20% Tris-glycine gel (Biorad). Blots were transferred to PVDF membrane (Biorad) and blocked with either 5% milk (Thermo Fisher Scientific) or 5% BSA (New England Biolabs) in phosphate buffered saline with 0.1% Tween (Boston Bioproducts, Ashland, MA), or Everyblot blocking buffer (Cat. No 12010020). Secondary detection of horseradish peroxidase-linked antibodies (Cell Signaling Technologies) with horseradish peroxidase substrate (Thermo Fisher Scientific) was visualized using Amersham ImageQuant 800 (Cytiva Life Sciences, Marlborough, MA). When needed, stripping was performed using the Restore™ PLUS Western Blot Stripping Buffer (Cat. No. 46430), and membrane was re-blocked with Everyblot blocking buffer.

Antibodies used as follows: 1:1000 dilution of anti-CHKA (Cell Signaling Technology (CST), Cat. No. 13422S) in Everyblot, 1:1000 anti-HDAC1 (CST, Cat. No. 5356S), 1:1000 anti-H4 (abcam, Cat. No. ab7311), 1:1000 anti-β-actin (CST, Cat. No. 4967L), secondary anti-rabbit IgG, HRP linked antibody (CST, Cat. No. 7074S, 1:1000), secondary anti-mouse, HRP linked antibody (CST, Cat. No. 7076S, 1:1000), 1:1000 anti-RAD51 (CST, Cat. No. 8875T), 1:1000 anti-GAPDH (CST, Cat. No. 2118S), 1:1000 anti-vinculin (CST, Cat. No. 13901S), 1:1000 anti-β-tubulin (CST, Cat. No. 2128S), 1:1000 anti-HA-tag (CST, Cat. No. 3724S), 1:1000 anti-MLX (CST, Cat. No. 85570S), 1:1000 anti-perilipin-2 (CST, Cat. No. 45535S), and 1:1000 anti-tom20 (CST, Cat. No. 42406S).

### Cell fractionation experiment

Five million Jurkat cells were pelleted by centrifugation at 300 g for 5 minutes at 4 °C. Cells were washed one time with 1 mL of ice-cold PBS and centrifuged at 300 g for 5 minutes at 4 °C. The cell pellet was resuspended in 250 µL of lysis buffer (10mM HEPES pH 7.5, 10mM KCl, 0.1mM EDTA, 1mM DTT, 0.5% NP-40, 1 X protease/phosphatase inhibitor cocktail) and incubated on ice for 20 min. 50 µL was collected as the whole cell lysate. The suspension was vortexed and then centrifuged at 12,000 g for 10 min at 4 °C. 200 µL of supernatant was collected as the cytoplasmic fraction. The nuclear pellet was resuspended in 200 µL of lysis buffer and spun at 200 g for 5 minutes at 4 °C for a total of four times. The nuclear pellet was resuspended in 250 µL of ice-cold fractionation buffer (2 M sucrose, 1 mM MgCl_2_, 10 mM Tris-HCl pH 7.4) and centrifuged at 16,000 g for 30 minutes at 4 °C. Nuclei were washed twice with 200 µL of lysis buffer at 200 g. The nuclei were resuspended in 100 µL of nuclear extraction buffer (20mM HEPES pH 7.5, 400 mM NaCl, 1 mM EDTA, 1X protease/phosphatase inhibitor cocktail) and incubated on ice for 5 minutes. The sample was spun at 16,000 g for 5 minutes.

The supernatant was collected as the nuclear fraction, and the pellet was resuspended in 4X Laemmli buffer + 4% BME as the chromatin fraction. CHKA was blotted for, and fractionation was confirmed via western blotting of control markers HDAC1, β-actin, and histone H4.

### RNA-sequencing

Total RNA was extracted from 1,000,000 Jurkat cells/sample using the RNeasy Mini Kit (QIAGEN, Cat. No. 74104) and 1 µL of 1:10,000 diluted ERCC RNA spike-in controls was added (Invitrogen, Cat. No. 4456740). Three replicates of each condition (shGFP, shLuciferase, shCHKA#3, and shCHKA#5) was performed. RNA was quantified using the Quant-iT RiboGreen RNA assay (ThermoFisher) and quality checked by the 2100 Bioanalyzer RNA 6000 Nano assay (Agilent) or 4200 TapeStation High Sensitivity RNA ScreenTape assay (Agilent) prior to library generation. Libraries were prepared from total RNA with the Illumina Stranded Total RNA Library Prep Kit according to the manufacturer’s instructions (Illumina, PN 20040529). Libraries were analyzed for insert size distribution using the 2100 BioAnalyzer High Sensitivity kit (Agilent), 4200 TapeStation D1000 ScreenTape assay (Agilent), or 5300 Fragment Analyzer NGS fragment kit (Agilent). Libraries were quantified using the Quant-iT PicoGreen ds DNA assay (ThermoFisher) or by low pass sequencing with a MiSeq nano kit (Illumina). Paired end 100 cycle sequencing was performed on a NovaSeq X Plus (Illumina).

Total stranded RNA sequencing data were pre-processed by the internal AutoMapper pipeline.^53^ Briefly the raw reads were first trimmed (Trim-Galore version 0.63), mapped to human (GRCh38) or mouse (mm10) genome assembly (STAR v2.79a)^54^ and then the gene level values were quantified (RSEM v1.3.3)^55^ based on GENCODE annotation (v31 for human, VM22 for mouse). Genes with low counts (counts per million mapped reads or CPM < 0.1) were removed from the analysis, and only protein-coding genes were used for differential expression analysis. Normalization factors were generated using the TMM method,^56^ counts were then transformed using voom^57^ and then analyzed using the lmFit and eBayes functions (R limma package version 3.42.2).^58^ The false discovery rate (FDR) was estimated using the Benjamini–Hochberg method. The cutoff for differentially expressed genes used was |log2FC|>0.5 & FDR < 0.05 and a p-value < 0.05.

### ATAC-sequencing

Accessible regions of DNA were extracted using the Active Motif^®^ ATAC-seq kit (Cat. No. 53150). Libraries were prepared according to the protocol. Briefly, a unique i7 and i5 index primer combination was added to the DNA, the product was PCR amplified, and SPRI clean-up was performed. Libraries were sequenced with Illumina Next Generation Sequencing. Peak annotation, differential peak calling, GO enrichment, and motif enrichment analysis were completed from the sequencing results. Gene set enrichment analysis was performed based on the following gene sets: hallmark, curated, transcription factor targets, computational, ontology and oncogenic signature gene sets. Three replicates of each condition (shGFP, shLuciferase, shCHKA#3, and shCHKA#5) were performed.

Briefly, raw reads in fastq format were processed with Trim-Galore tool^59^ to remove potential adapters and quality trim 3’ end of reads with cutadapt program,^60^ followed by FastQC analysis.^61^ A quality score cutoff of Q20 was used and the first 15bp of each read were also clipped to reduce the GC bias. Next, reads were mapped to human reference genome (hg38; GRCh38.p12) with BWA mem (0.7.17-r1188),^62^ then converted to BAM format and deduplicated with fq2bam (v3.0.0.6). Subsequently, uniquely mapped properly paired reads were extracted with samtools^63^ (v1.2) and fragments were extracted with bedtools^64^ (v2.24.0).

### Homologous recombination assay

100,000 Jurkat cells per well were transfected with 0.5 µg of each dl plasmid (dl-1 and dl-2) from the Norgen Biotek Homologous Recombination Assay Kit (Cat. No. 35600). For the positive control, 25 ng of the positive control plasmid was transfected. For the negative control, 25 ng of each of the dl plasmids was used in separate transfections. After 48 hours, the total DNA was isolated using the PureLink® Genomic DNA Kit (Cat. No. K1820-01). 20µL PCR reactions were set up in quadruplicates using the Luna^®^ Universal qPCR Master Mix (Cat. No. M3003S), 2 µl of the provided assay primers mixture, and 50 ng of plasmid DNA per sample. To detect the backbone plasmids, the same PCR reactions was set up using the supplied universal primers mixture instead. qPCR was performed using the following protocol: Initial denaturation at 95°C for 3 minutes then 95°C for 15 sec, 61°C for 15 sec, and 72°C for 15 sec, and repeated for 40 cycles then run the melt curve analysis. Samples were normalized with universal primers samples. The amount of recombinant product for each reaction was using the ΔΔCt method against the negative control.

### Immunofluorescence

500,000 shLuciferase or shCHKA cells were treated with 1mM of oleic acid-albumin from bovine serum (Cat. No. O3008) or albumin from bovine serum for 24 hrs prior to the immunofluorescence experiment. No oleic acid was added to the cells in Figure 4A. The cells were harvested and suspended in 1X PBS. 200,000 cells/slide were spun at 600 rpm for 5 minutes using the ThermoScientific cytospin 4 instrument to poly-lysine microscope slides (Cat. No. P0425). Slides were dried at room temperature for 5 minutes and fixed in 4% paraformaldehyde in PBS for 15 minutes. Slides were washed 3X with 1X PBS for 5 minutes each. Cells were incubated with 2µM BODIPY 493/503 in PBS for 30 minutes at room temperature covered from light. Cells were washed one time with 1X PBS for 5 minutes. ProLong™ Diamond Antifade Mountant with DAPI (Cat. No. P36971) was used to mount coverslips to the slides. The coverslips were sealed with clear nail polish and dried overnight protected from light. Cells were imaged with either a Zeiss 780 microscope with the 63X objective or performed on a custom-built Nikon microscope equipped with a Yokogawa CSU-W1 Spinning Disk unit, Z-axis piezo nanopositioners, and Tokai Hit STXG CO2 incubation system using a 60x (CFI APO TIRF, NA 1.49 oil) objective. The surface area of lipid droplets/cell was counted from 50 cells from each condition. Images were processed using FIJI (2.16.0). Brightness/contrast was adjusted equally across shLuciferase and shCHKA images from each experiment to ensure valid comparison. Raw images in Figure 4A were cropped using the “rectangle” tool to ensure equal cropping of both images at the dimensions of width: 128.6012 microns and height: 123.4443 microns.

### Lipid droplet extraction

Lipid droplets were extracted from 15,000,000 shLuciferase or shCHKA Jurkat cells using the Lipid Droplet Isolation Kit (abcam, Cat. No. ab242290). Samples were tested for the presence of different proteins (see western blot section).

### Statistical Analyses

For two-group comparisons of continuous measures, a two-tailed Welch unequal variances t-test was used, except otherwise stated. For 3-group comparisons of continuous measures, a one-way analysis of variance model (ANOVA) was performed and a Dunnett correction for multiple comparisons was used. For analysis of two effects, a two-way ANOVA model was constructed with either Dunnet, Sidak, or Tukey’s adjustments for multiple comparisons, as appropriate.

## DATA AVAILABILITY

The RNA and ATAC sequencing data generated in this study have been deposited in the NCBI Gene Expression Omnibus (GEO) database under accession code GSE350591 (RNA-seq) and GSE350590 (ATAC-seq). These data will be made public concurrent with publication of this manuscript and are available to the reviewers by searching for the above accession numbers at https://www.ncbi.nlm.nih.gov/geo/, using the following reviewer tokens: Reviewer token for GSE350591: snqpggwgdzqnfkj. Reviewer token for GSE350590: ypgzgswefvmbxox.

All raw and annotated immunofluorescence and western blot images have been uploaded to Mendeley and will be made public concurrent with publication. Prior to publication, these data are available to the reviewers at https://data.mendeley.com/preview/b8sfzptwm9?a=194381bf-e95a-42ea-b50b-4e8c2c29cd36. Any other material is available upon request.

## ACKNOWLEDGEMENTS

We are grateful to all members of the A.G. laboratory for advice and recommendations, and Kathryn E. Wellen and Christina Demetriadou for experimental advice. This work was supported by NIH grants CA193651 and CA316383 and Alex’s Lemonade Stand Foundation for Childhood Cancer.

