## Supplemental Data Figures and Legends for "Choline kinase A is required for alkylating agent resistance via lipid droplet formation"

[1]Department of Oncology, St. Jude Children's Research Hospital, Memphis, TN 38105, USA.
[2]St. Jude Graduate School of Biomedical Sciences, Memphis, TN 38105, USA.
[3]Division of Hematology/Oncology, Boston Children's Hospital, Boston, MA, 02115, USA.
[4]Department of Cell & Molecular Biology, St. Jude Children's Research Hospital, Memphis, TN 38105, USA.
[5]Center for Applied Bioinformatics, St. Jude Children's Research Hospital, Memphis, TN 38105, USA.
[6]Department of Surgery and Cancer, Faculty of Medicine, Imperial College London, London SW7 2BX, UK.
[7]Department of Biochemistry and Molecular Genetics, University of Colorado Anschutz, Aurora, CO 80045.
[8]Department of Pediatric Oncology, Dana-Farber Cancer Institute, Boston, MA, 02215, USA.
*Equal contribution

**Figure S1**

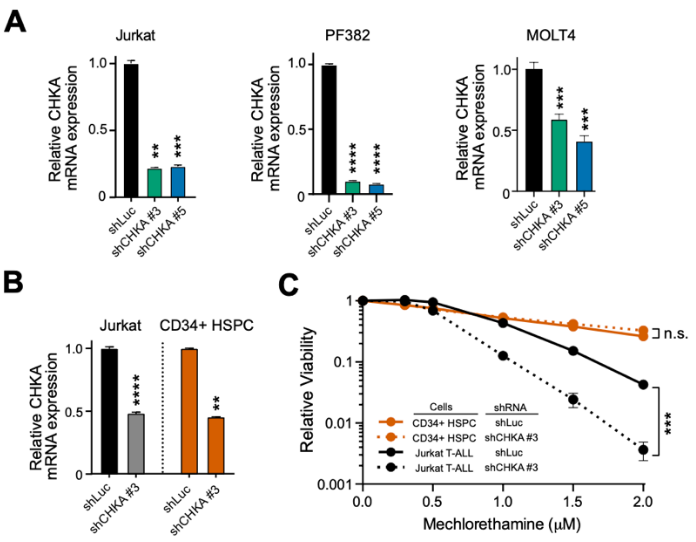


**Figure S1. Effects of CHKA knockdown in leukemia and normal cells.** (A) The indicated T-ALL cells were transduced with CHKA or control (shLuciferase)-targeting shRNAs, and knockdown efficiency was assessed by quantitative RT-PCR analysis, with β-actin (*ACTB*) as the control. Significance was assessed by 2-way ANOVA with Dunnett adjustment for multiple comparisons. (B) The indicated cell lines were transduced with control (shLuc) or CHKA-targeting shRNAs, and knockdown efficiency was assessed by quantitative RT-PCR analysis. Significance was assessed by Welch's t test within each cell type. (C) Cells from (B) were treated with mechlorethamine at the indicated doses for 5 days, and viability was assessed from live cells counts after trypan blue vital dye staining. All cell counts are normalized to those in shLuciferase-transduced, no-drug controls. Significance was assessed by 2-way ANOVA with Sidak adjustment for multiple comparisons at the highest mechlorethamine dose shown. All data points represent the mean ± SEM of at least n = 3 biological replicates.

*, $p < 0.05$; **, $p < 0.01$; ***, $p < 0.001$; ****, $p < 0.0001$; n.s., $p \geq 0.05$.

**Figure S2**

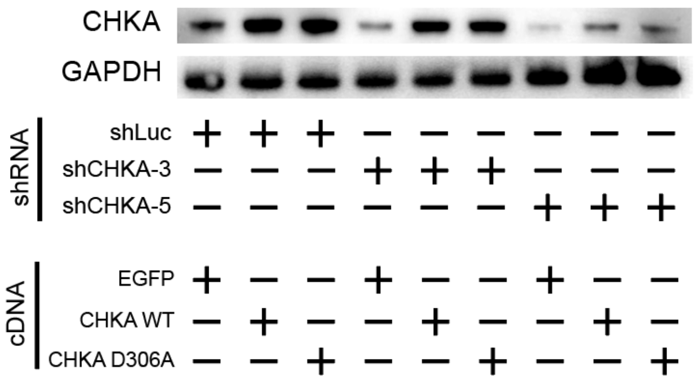


**Figure S2. Western blot analysis for rescue of CHKA-targeting shRNAs by cDNA rescue constructs.** Jurkat cells were transduced with the indicated shRNA and cDNA expression constructs, and expression of the indicated proteins was assessed by Western blot analysis.

### Figure S3

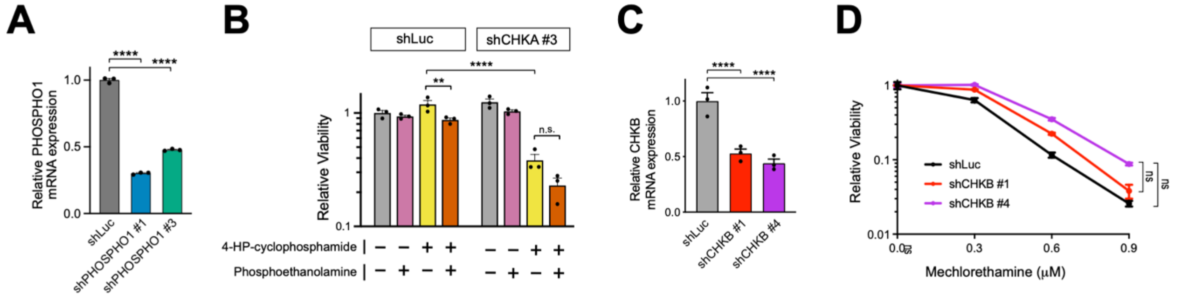


**Figure S3. Effects of CHKB and PHOSPHO1 knockdown on mechlorethamine response.** (A) Jurkat cells were transduced with the indicated shRNAs, and knockdown efficiency was assessed by quantitative RT-PCR analysis, with β-actin as the control. Significance was assessed by one-way ANOVA with Dunnett adjustment for multiple comparisons. (B) Jurkat cells were transduced with the indicated shRNAs and supplemented with vehicle or phosphoethanolamine (1 mM added daily) beginning 24 hours prior to starting treatment with 4-HP-cyclophosphamide. Relative cell viability was assessed after 5 days of treatment. A two-way ANOVA with Sidak adjustment for multiple comparisons was used for statistical analysis. (C) Jurkat cells were transduced with control or CHKB-targeting shRNAs, and knockdown efficiency was validated by quantitative RT-PCR analysis. Significance was assessed by one-way ANOVA with Dunnett adjustment for multiple comparisons. (D) Cells from (C) were treated with mechlorethamine at the indicated doses for 5 days, and viability was assessed. Significance was assessed by Welch's t test.

*, $p < 0.05$; **, $p < 0.01$; ***, $p < 0.001$; ****, $p < 0.0001$; n.s., $p \geq 0.05$.

**Figure S4**

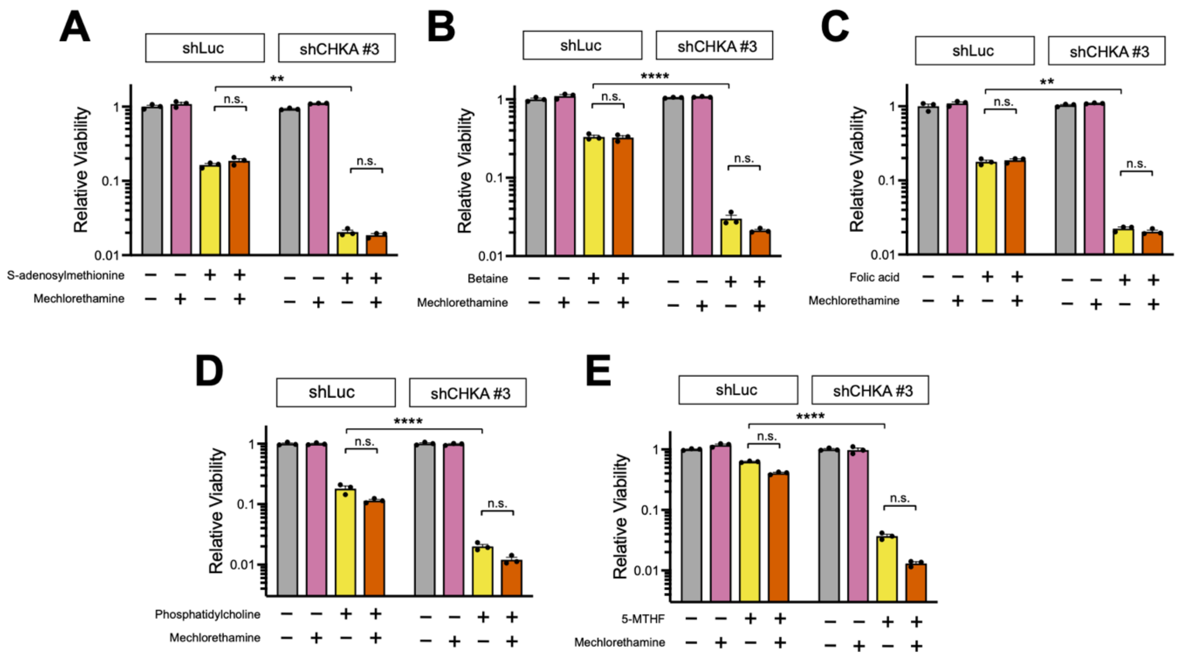


**Figure S4. Supplementation of additional choline-related metabolites fails to protect CHKA-deficient cells from the toxicity of alkylating agents.** (A-E) Jurkat cells were transduced with the indicated shRNAs, treated with the indicated molecules, and viability was assessed by Cell TiterGlo analysis after 5 days of treatment with 1.5 μM 4-HP-cyclophosphamide. Doses used were: S-adenosylmethionine, 50 μM; mechlorethamine, 1.5 μM; betaine, 5 mM; folic acid, 100 μM; phosphatidylcholine, 100 μM; 5-methyltetrahydrofolate (5-MTHF), 150 μM. Significance was assessed by 2-way ANOVA with Sidak adjustment for multiple comparisons. All data points represent the mean +/- SEM of at least 3 biological replicates.

*, $p < 0.05$; **, $p < 0.01$; ***, $p < 0.001$; ****, $p < 0.0001$; n.s., $p \geq 0.05$.

**Figure S5**

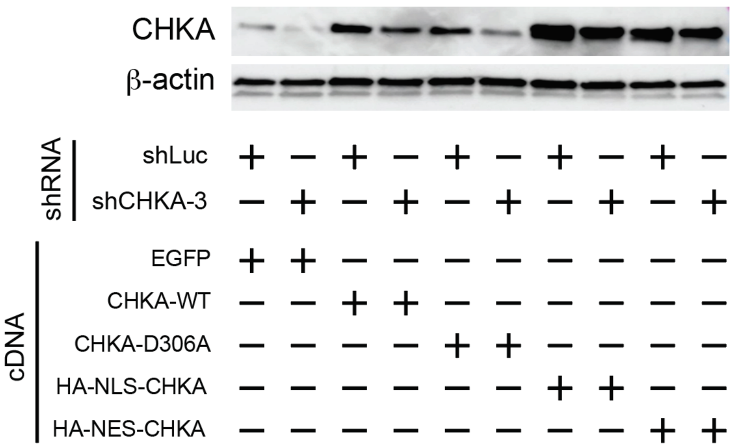


**Figure S5. Effects of shRNA knockdown on CHKA protein levels and their rescue by cDNA rescue constructs utilized.** Jurkat cells were transduced with the indicated shRNA and cDNA expression constructs, and expression of the indicated proteins was assessed by Western blot analysis.

**Figure S6**

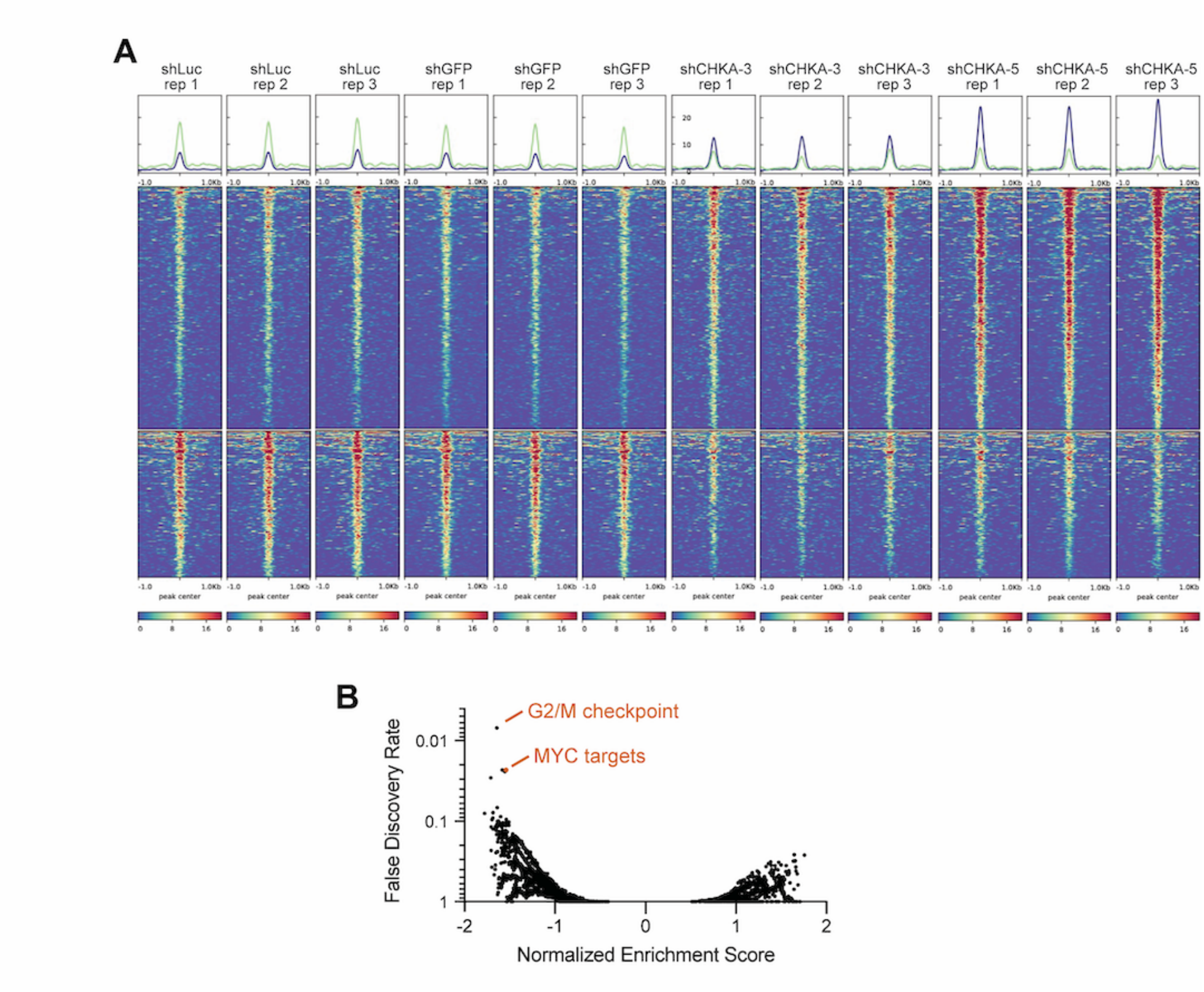


**Figure S6. CHKA knockdown alters chromatin accessibility.** (A) Jurkat cells were transduced with the indicated shRNAs, and ATAC-seq analysis was performed to map chromatin accessibility across the genome. Genomic regions whose accessibility changed most are shown. (B) Gene set enrichment analysis of ATAC-seq peaks that changed in accessibility following CHKA knockdown, with MYC target genes among the most downregulated.

**Figure S7**

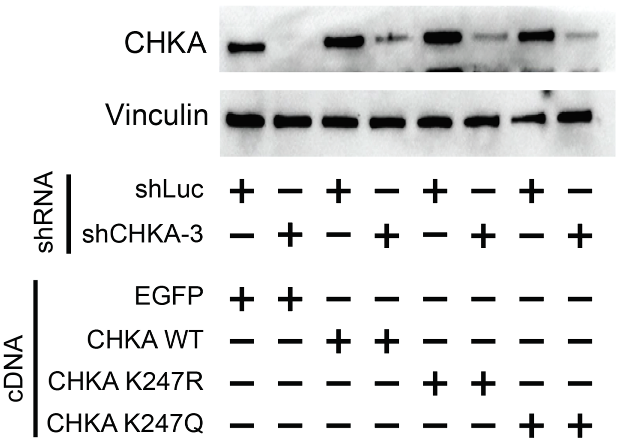


**Figure S7. Effects of CHKA knockdown and rescue on expression of the protein for CHKA K247R/Q mutants.** Jurkat cells were transduced with the indicated shRNA and cDNA expression constructs, and expression of the indicated proteins was assessed by Western blot analysis.
